# Ribosomes as molecular thermometers: metal-binding sites in ribosomal proteins are robust indicators of bacterial adaptation to heat and cold

**DOI:** 10.1101/2022.09.09.507293

**Authors:** Antonia van den Elzen, Karla Helena-Bueno, Charlotte R. Brown, Sergey Melnikov

## Abstract

Ribosomal genes are widely used as “molecular clocks” to infer the evolutionary relatedness of species. It is unclear, however, whether these genes can also serve as “molecular thermometers” to precisely estimate an organism’s optimal growth temperature. Previously, some estimations were made using the average nucleotide content in ribosomal RNA, but the universal application of this approach was prevented by numerous outliers. Here, seeking to bypass this problem, we asked whether ribosomal genes contain additional markers of thermal adaptations, aside from their nucleotide composition. To answer this, we analyzed site-specific variations in sequences of ribosomal proteins from 2,021 bacteria with known optimal growth conditions. We found that ribosomal proteins comprise a few “mutational hotspots”—residues that vary in a temperature-dependent manner and distinguish heat- and cold-adapted bacteria. Most of these residues coordinate metal ions that support protein folding at high temperatures. Using these residues, we then showed that the upper and lower limits of an organism’s optimal growth temperatures can be estimated using just 0.001% of the genome sequence or just two amino residues in the cellular proteome. This finding illustrates that laboratory-independent estimation of optimal growth temperatures can be simplified if we abandon the traditional use of rRNA and protein sequences to assess their content and instead focus on those few residues that are most critical for protein structure. This finding may simplify the analysis of unculturable and extinct species by helping bypass the need for laborious, costly, and at times impossible laboratory experiments.

## Introduction

Genes coding for ribosomal RNA and ribosomal proteins are widely used as molecular clocks to infer phylogenetic relationships among species and gain insights into the origin of life^1-12^. It is unclear, however, if these genes contain other useful information, such as information about the optimal growth environment for a given species.

Previous studies found that ribosomal genes contain some information about a given organism’s optimal growth temperature^13-23^. For instance, thermophilic ribosomal proteins were found to have a higher content of arginine, isoleucine, proline, and tyrosine, and a decreased content of serine and threonine. However, this correlation was not consistent across species to serve as an accurate predictor of an organism’s optimal growth temperature^13,14^. A more promising approach was established by studying the correlation between the optimal growth temperature and the G+C content in rRNA^15-23^. For example, an almost linear correlation was observed in 143 microbial genera, where the rRNA G+C content gradually increased from about 45% G+C in psychrophilic bacteria to about 55% G+C in thermophilic bacteria^15^. But at least 22 other studied genera showed temperature-independent variations because the rRNA G+C content was found to be influenced by other factors, including high-salt conditions and host-restricted lifestyles^15,24-29^. Collectively, these studies showed that using the content of rRNA genes as a molecular thermometer is a promising approach but on its own it lacks accuracy, raising the question: Do ribosomal genes contain other markers of thermal adaptation that could help estimate the optimal growth conditions for a given organism?

Here, we addressed this question by correlating the experimentally defined growth temperatures for 2,021 representative bacteria with the sequences of their ribosomal proteins. Our goal was to abandon the traditional approach in which sequences of rRNA or proteins are treated as strings of equally important residues to calculate their contents. Instead, we focused our attention on the very small number of residues that were shown to be critically important for the stability of the folding of ribosomal proteins in some thermophiles. Specifically, studies of *Thermus thermophilus* ribosomes revealed that the smallest ribosomal proteins coordinate metal ions to enable protein folding at high temperatures^30-34^. The protein uS4 coordinates an iron-sulfur cluster^30^ and the proteins uS14, bS18, uL24, bL28, bL31, bL32, bL33, and bL36 coordinate zinc ions^31-34^.

Given this stabilizing impact of the metal coordination, we hypothesized that the 36 metal-binding residues in ribosomal proteins can indicate the optimal growth conditions of a living cell. We then tested this hypothesis using the analysis of over 24,000 sequences of ribosomal proteins. We found that the occurrence of metal-binding residues does indeed correlate with the optimal growth temperature. We finally showed how to use this finding to accurately estimate the upper or lower limits of optimal growth temperature based on a sequence of just two amino acid residues in a cellular proteome. Overall, our work illustrates that studying site-specific variations—as opposed to variations in content—can reveal new robust markers of environmental adaptation in the most conserved genes in nature. These newly identified markers provide an independent tool to study the ancient history of climate change by sequencing the ribosomal genes of extant and extinct species.

## Results

### Metal-binding residues are widespread in ribosomal proteins from heat-adapted bacteria

We first asked whether metal-binding residues that were found in ribosomal proteins from *Thermus thermophilus* are present in other heat-adapted species. To test this, we retrieved 24,312 sequences of the ribosomal proteins the uS4, uS14, bS18, uL24, bL28, bL31, bL32, bL33, and bL36 from 2,021 bacteria (from 825 genera) using our recently established database of organisms with experimentally defined optimal growth temperatures (**Table S1, Methods**)^35^. We focused on bacteria because they exist in thermal equilibrium with their environment, and the genome sequences are available for many species with identified optimal growth temperatures. We then aligned these protein sequences and assessed how the identity of each residue in each protein depends on optimal growth temperature (**Fig. 1, Fig. S1**).

**Figure 1.**
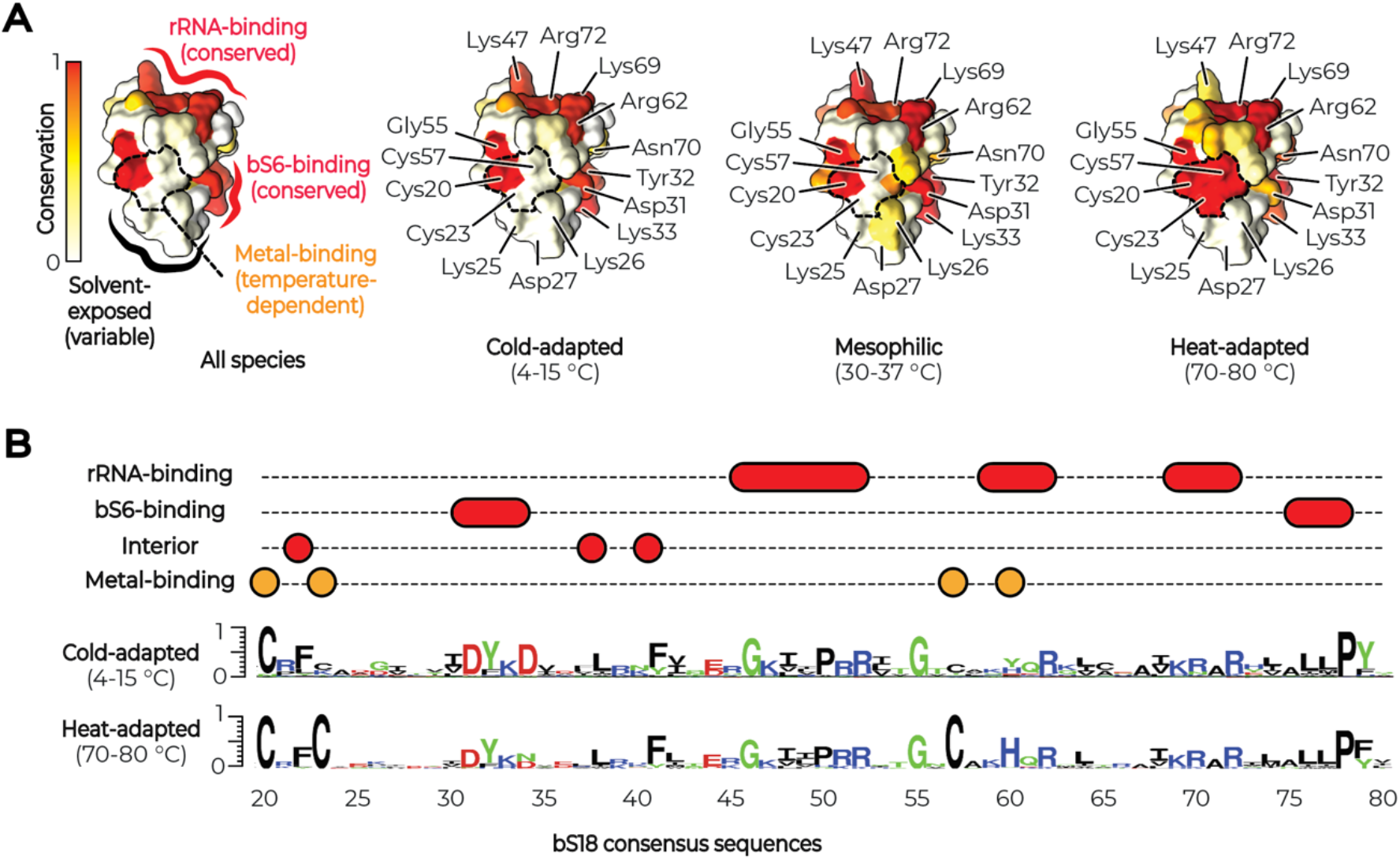
Metal-binding residues are widespread in ribosomal proteins from heat-adapted bacteria. (**A**) The structure of the *E. coli* ribosomal protein bS18 shows how the conservation of protein residues depends on the optimal growth temperature. bS18 possesses three groups of residues. The first group includes residues that remain nearly invariant across all bacteria, regardless of their optimal growth temperature. Most of these mediate bS18 binding to the ribosome. The second group includes residues that remain highly variable across all bacteria, regardless of their optimal growth temperature. These residues are predominantly found on the solvent-side of a protein. Only the third and smallest group includes residues whose conservation depends on the optimal growth temperature of bacterial species. These residues remain highly conserved in heat-adapted bacteria but become highly variable in cold-adapted species. In the bS18 structure, these residues coordinate Zn^2+^ ions, which appears to stabilize bS18 folding at high temperatures. (**B**) Conservation “logos” illustrate that the metal-binding residues (Cys23, Cys55, and His60) are the only conserved bS18 residues that distinguish heat-adapted species from non-heat-adapted species. Thus, out of 75 amino acid positions in the structure of bS18, only three positions show robust changes in amino acid sequence in a temperature-dependent manner.

This analysis revealed that non-thermophilic species possess just a small fraction of highly conserved residues (**Fig. 1, Fig. S1**). For example, protein bS18 comprises just 21 conserved residues that mediate bS18 binding to the ribosome and support its folding. In thermophiles, we found the same 21 conserved residues plus three additional conserved residues—Cys23, Cys55, and His60—that stabilize bS18 folding by coordinating a Zn^2+^ ion. Similarly, proteins uS4, uS14, uL24, bL28, bL31, bL32, bL33, and bL36 possess just 45 residues that are highly conserved in thermophiles but not in non-thermophilic species (**Fig. 1, Fig. S1**). Of these residues, 33 coordinate Zn^2+^ ions (in uS14, uL24, bL28, bL31, bL32, bL33, and bL36) and an iron–sulfur cluster (in uS4). Thus, the metal-binding residues in ribosomal proteins are indeed widespread among thermophilic species, and they represent a very small fraction of residues that distinguish all lineages of thermophilic bacteria from non-thermophilic bacteria.

### Metal-binding residues indicate the limits of the optimal growth temperature

We next asked if the occurrence of the metal-binding residues correlate with the optimal growth temperature. And, if so, can we use these residues to estimate the optimal temperature for bacterial growth. To test this, we pooled the sequences of ribosomal proteins into eight bins, corresponding to optimal growth temperature intervals from 1–10°C to 81–90°C. Then, for each ribosomal protein, we calculated the frequencies of the metal-binding ribosomal proteins in each bin (**Methods**).

We found that each ribosomal protein gradually “transforms” from a metal-free isoform (C-isoform) to a metal-binding isoform (C+ isoform) upon transition from cold-adapted to heat-adapted species (**Fig. 2**). For example, in protein bL32, the metal-binding residues are strictly absent in organisms with optimal growth temperatures between 4 and 20°C. Between 21 and 60°C, these residues become more prevalent, and above 60°C, they become strictly conserved.

**Figure 2.**
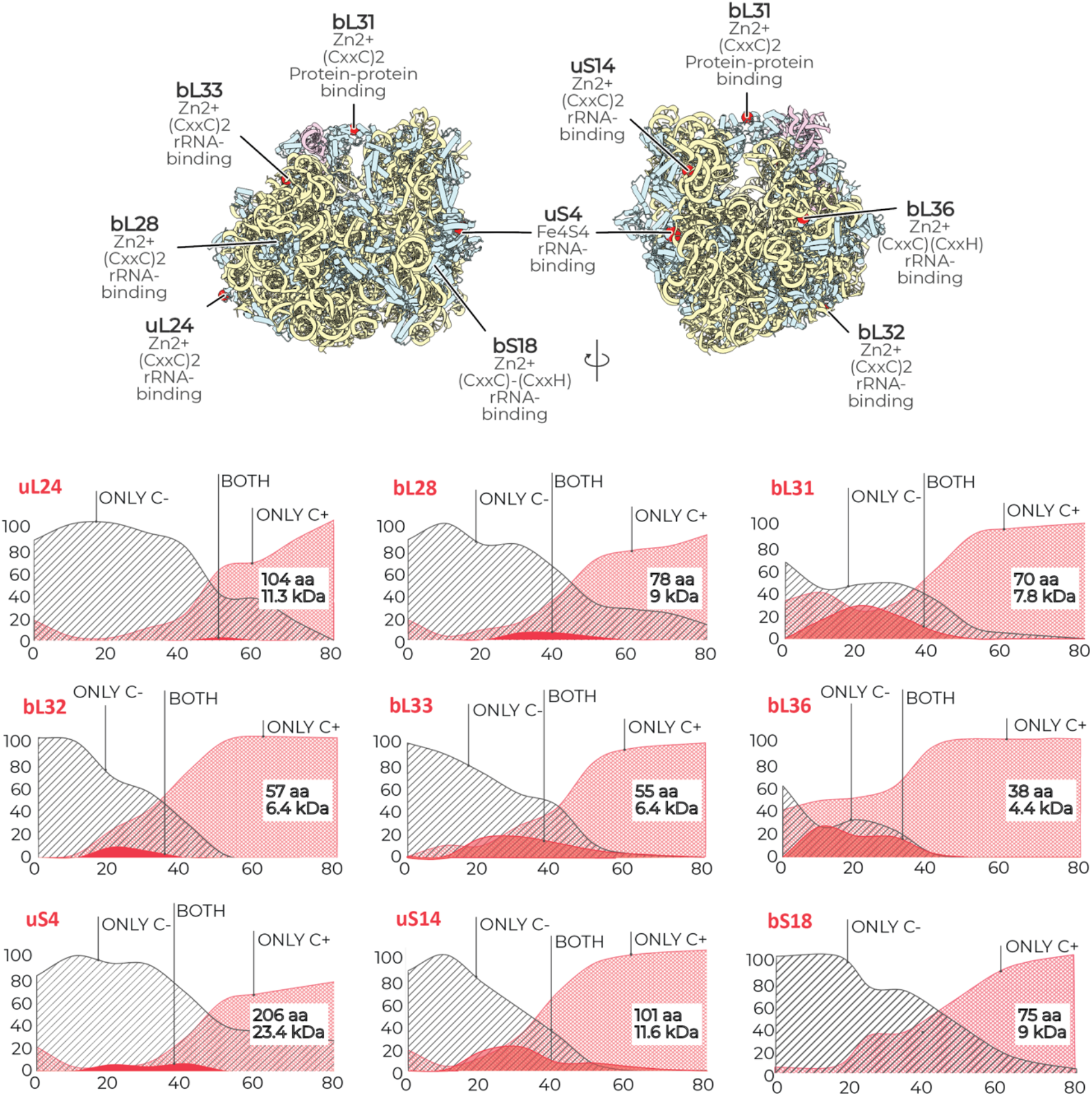
Metal-binding residues indicate the limits of the optimal growth temperature. The structure of bacterial ribosomes from *T. thermophilus* (pdb id 4y4o) illustrating the location of metal-binding sites (CxxC or CxxH) in ribosomal proteins, including eight proteins that bind Zn^2+^ ions and one that binds an iron–sulfur cluster (Fe_4_S_4_). The plots show how the occurrence of metal-binding sites in each of the ribosomal proteins changes with the optimal growth temperatures of bacterial species: upon transition from thermophiles to psychrophiles, the occurrence of metal-binding sites decreases in each ribosomal protein.

We also found that the conservation of the metal-binding residues depends on the protein size (**Fig. 2**). For example, in the smallest ribosomal protein (bL36), the metal-binding sites are immutable in all bacteria with optimal growth temperatures of 40°C or more. In larger proteins (uS14, bL31, bL32, and bL33), the metal-binding sites are immutable in bacteria with optimal growth temperatures of 60°C or more. And in even larger proteins (uL24), the metal-binding sites are immutable in bacteria with optimal growth temperatures of 80°C or more. Thus, we found the metal-binding site (or their absence) may indicate the lower and upper limits of the optimal growth temperature for bacterial species.

At lower temperatures, however, we observed a few “outliers”—cold-adapted organisms that bear C+ ribosomal proteins (**Fig. 2, Supplementary Data 2**). We therefore decided to explore the nature of these “outliers”.

### Metal-binding residues are also conserved in ancient branches of non-thermophilic bacteria

We first asked if these outliers belong to the same branch of bacteria. To answer this, we calculated the total number of the metal-binding ribosomal proteins (C+ proteins) in each species to identify every species with “anomalously” high or low numbers of C+ proteins. We found that, on average, the number of C+ ribosomal proteins gradually increases from psychrophiles (1.5) to thermophiles (8.1) (**Fig. 3A, Table S1**). But 254 bacteria deviated from this tendency by having at least three more or less C+ proteins compared with their average number in a given temperature interval (**Fig. 3B, Supplementary Data 2**).

**Figure 3.**
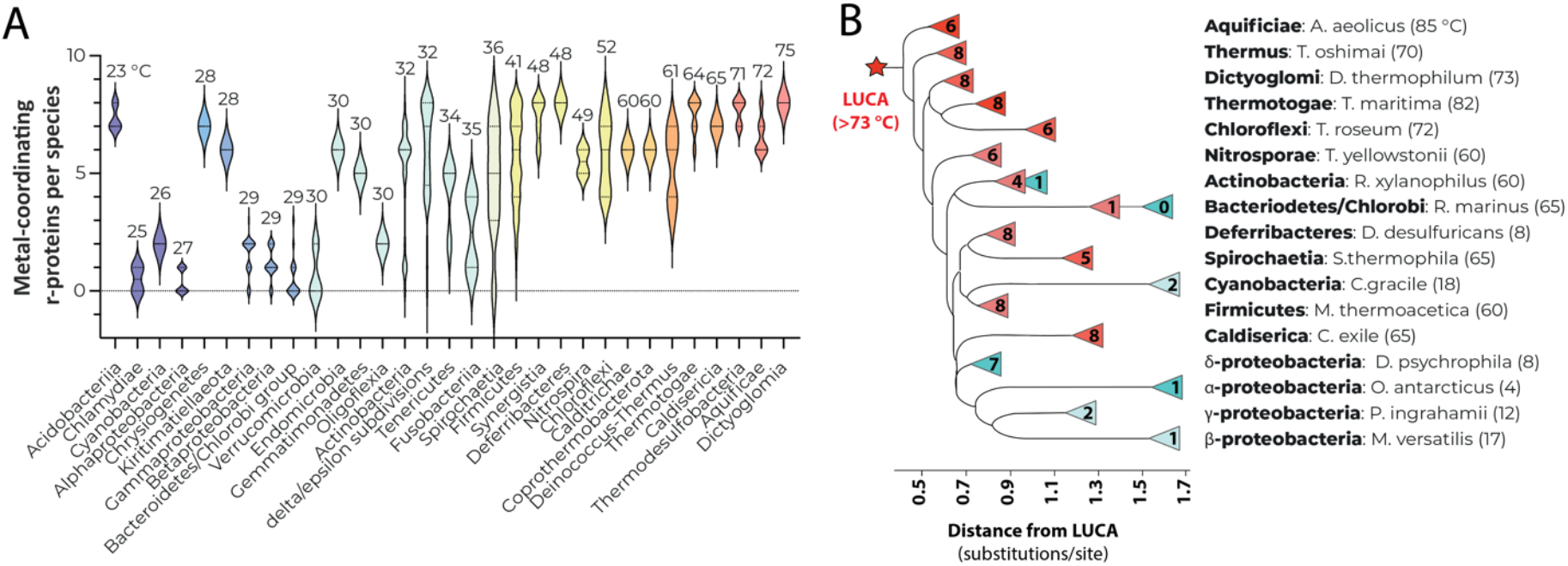
Metal-binding residues are also conserved in ancient branches of non-thermophilic bacteria. (**A**) Bacterial phyla are shown in the order of their ascending average optimal growth temperature to illustrate a gradually increasing number of metal-binding ribosomal proteins upon transition from predominantly cold-adapted to predominantly heat-adapted species. This panel also illustrates a few exceptions from this general tendency, such as the phyla of *Acidobacteria, Chrysiogenetes* and *Kiritimatiellaeota*. (**B**) A schematic tree of life shows that non-thermophilic species with high number of metal-binding ribosomal proteins tend to be located closer to the root of the tree of life. However, the “outliers” (species with “anomalously” few or many C+ ribosomal proteins) do not follow this pattern. For example, δ-proteobacteria possess metal-binding sites in most ribosomal proteins despite being psychrophiles or mesophiles, and they are located as close to LUCA as most thermophiles. Similarly, the heat-adapted bacteria *Rhodotermus marinus* has an optimal growth temperature of 65°C but is atypically remote from the root of the tree compared with other thermophiles and has only one C+ ribosomal protein. These outliers show that the distance from the root of the tree of life correlates with the anomalously high number of metal-binding sites in some cold-adapted bacteria and their anomalously low number in some heat-adapted bacteria.

We then mapped these outliers on the tree of life and found that they belong to different bacterial branches, indicating that C+ ribosomal proteins occasionally occur in evolutionarily distant non-thermophiles (**Fig. 3A,B**, **Supplementary Data 2**). Although the outliers belong to different branches of the tree of life, mapping them on the tree of life revealed one common property: their anomalous proximity to LUCA (**Fig. 3B**). Specifically, we found that thermophiles tend to stay closer to LUCA (0.91 substitutions per site in ribosomal proteins) compared to psychrophiles (1.29 substitutions per site), which is consistent with the current theory that life was born in hot environments, with the psychrophiles emerging when the Earth became cooler^36^. However, the outliers deviate from this evolutionary tendency (**Fig. 3B, Supplementary Data 2**). For example, the δ-proteobacteria *Desulfotalea psychrophila* has an optimal growth temperature of 8°C but bears 8 C+ ribosomal proteins. On the tree of life, this bacterium is located as close to LUCA as an average thermophile (**Fig. 3B**). Overall, this anomalous proximity to LUCA suggests that most outliers could emerge relatively early in the evolution, when the Ocean’s temperature was still relatively high.

## Discussion

### Ribosomal genes possess site-specific markers of thermal adaptation

In this study, we assessed the potential of using site-specific variations in metal-binding ribosomal proteins to estimate the optimal growth temperature of bacterial species. We found that sequences of ribosomal proteins contain a small fraction of residues that distinguish thermophilic organisms from non-thermophilic organisms. In doing so, we have identified a previously unknown class of site-specific markers of thermal adaptation in ribosomal genes, providing a tool to infer the limits of optimal growth conditions through sequencing of nature’s most common genes.

Aside from this fundamental finding, our study describes two instrumental advances. Firstly, by identifying mutational hot-spots in ribosomal genes, we show how to drastically reduce the amount of data required for laboratory-independent estimation of optimal growth conditions. Secondly, we provide the first ever analysis of site-specific variations in ribosomal genes in organisms with differing optimal growth temperatures. This analysis illustrates that studying site-specific variations—as opposed to variations in content—can reveal previously unknown and robust markers of thermal adaptation, which have remained elusive for more than three decades of conventional studies of ribosomal genes. Our analysis shows that only 1 in about 25 residues in ribosomal proteins shows lineage-independent differences between heat- and cold-adapted bacteria. This explains why the metal-binding residues were not identified earlier as markers of thermal adaptation: they are present in just 9 out of 54 ribosomal proteins and account for just 0.54% of residues in ribosomal proteins. Yet, the important location of these residues in protein structures appears to create a particularly strong selective pressure to retain these residues in heat-adapted species, rendering them markers of thermal adaptation.

### How can we use metal-binding residues as molecular thermometers?

The occurrence of metal-binding sites is binary: they are either present in proteins or not. Hence, each of these residues in isolation cannot be used as a thermometer to accurately predict the value of the optimal growth temperature for a given species. However, because different proteins have different temperature thresholds at which these residues become immutable, we can use these residues to estimate the upper or lower limits of the optimal temperature.

For example, if we find an organism that bears the metal-binding residues in bL32, then its optimal growth temperature is likely to exceed 20*°C—*because the metal-binding residues in bL32 are strictly absent in bacteria adapted to growth below 20°C. And if this organism lacks the metal-binding residues in bL36, then its optimal growth temperature is likely to be less than 40°C—because the metal-binding residues in bL36 are strictly conserved in organisms adapted to growth above 40°C. Thus, just eight residues in the cellular proteome—which correspond to approximately 0.004% of the average bacterial genome—appear to provide robust information about the optimal growth temperatures of bacterial species. What’s more, typically ribosomal proteins have either all four metal-coordinating residues (in thermophiles) or none of them (in non-thermophiles), which makes it possible to reduce the number of analyzed metal-coordinating residues by four. Thus, a few specific sites in the most widespread genes in nature can be used to estimate the optimal growth temperature of difficult-to-grow or extinct bacteria.

### What is the basis for using ribosomal genes as molecular thermometers?

Ribosomal genes are widely used to study the origin and evolution of life on our planet. These genes are estimated to originate about 3.9 billion years ago, and they are found in every organism known to science, including extinct species^37^. Yet we still do not know how much information about the origin and evolution of life is written in sequences of these ancient and ubiquitous genes.

Ribosomal genes are used as molecular clocks because the longer organisms exist as evolutionary distinct species, the more diverse their ribosomal genes become. But what is the basis for using ribosomal genes as molecular thermometers?

Previous studies suggest this basis may stem from the environmental history of life on Earth. Life on Earth is thought to have emerged in hot environments, with LUCA being either a thermophile or a hyperthermophile^36^. Specifically, the analyses of minerals^38-40^ and resurrected enzymes^41-43^ suggests that Earth’s surface has gradually cooled down from 75°C about 3 billion years ago, to 35°C about 420 million years ago, with a further decrease to 14°C today. It is therefore likely that—if the metal-binding residues pre-existed in LUCA due to their requirement for protein folding at high temperatures—organisms invading colder environments could gradually mutate these residues to endow proteins with the flexibility required in colder environments^44-46^.

Then why do the metal-binding residues are also present in some cold-adapted species? The answer to this question is unclear but we see three possible reasons. Firstly, the evolution of the metal-binding residues can be influenced by factors other than temperature. Secondly, this may stem from the fact that different branches of cold-adapted bacteria have likely emerged independently from each other and at different points of time. For example, previous reconstructions of Earth’s environment suggested that such “outliers” as δ-proteobacteria have emerged when the oceans temperature varied between 55 and 65°C, whereas other branches of proteobacteria have emerged when the oceans temperature dropped below 39°C^41^. Thirdly, the occasional occurrence of metal-binding residues in non-thermophilic species could reflect alternative mechanisms of cold adaptation. For example, previous studies showed that marked leaps in cold tolerance can be achieved by improving the process of protein folding: the expression of the cold-adapted chaperonins GroEL and GroES enabled *E. coli* growth at -13.7 °C, rendering *E. coli* a cold-tolerant organism without a single mutation in *E. coli* protein sequences^47^.

## Methods

### Annotating bacterial species with their optimal growth temperatures

The values of experimentally determined optimal growth temperatures were scraped from 23 public repositories of microorganisms, including the ATCC, the DSMZ and others (**Table S1**). The retrieved values were then added to the list of representative bacteria with fully sequenced genomes (https://www.ncbi.nlm.nih.gov/genome/browse). The resulting file (**Supplementary Data 1**) was then deposited in the database of organisms with experimentally defined optimal growth temperatures (http://melnikovlab.com/gshc/).

### Retrieving sequences of bacterial ribosomal proteins

To retrieve sequences of ribosomal proteins, we searched the Uniprot databank for protein sequences that are named “30S ribosomal protein X” or “50S ribosomal protein Y”, where X and Y corresponded to the bacterial name of ribosomal proteins from the small and large subunits, respectively. These data were then cleaned from truncated sequences by removing those sequences that were at least 25% shorter than the average length of a given protein in our dataset. The resulting files contained ∼2,400 sequences per each ribosomal protein (**Supplementary Data 2**).

### Assessing temperature-associated variations in bacterial ribosomal proteins

To identify temperature-associated variations in the ribosomal proteins, we aligned protein sequences using Clustal Omega^48^ with default settings (**Supplementary Data 3**), and then analyzed these alignments using the BioAlign package from BioPython^49^. In this analysis, for each ribosomal protein, we calculated two consensus sequences: a consensus sequence for cold-adapted bacteria (adapted to growth below 20°C) and a consensus sequence for heat-adapted bacteria (adapted to growth above 60°C). We then compared these consensus sequences using two different strategies. In the comparison strategy, we identified those residues that are highly conserved in both consensus sequences (more than 60% conserved) but have different identity. For example, this search would reveal a scenario in which a ribosomal protein carries a highly conserved phenylalanine residue in cold-adapted bacteria that is mutated to a highly conserved tyrosine in heat-adapted bacteria. In the second search, we identified those residues that change their conservation in heat-adapted bacteria compared to cold-adapted bacteria (using the 60%). This search would identify a scenario in which a ribosomal protein would have a poorly conserved residue in cold-adapted bacteria (e.g. present in less than 40% of sequences) but becomes highly conserved (e.g. 100% conserved) in heat-adapted bacteria. The resulting findings of this analysis were combined and mapped on the three-dimensional structures of ribosomal proteins using ChimeraX^50^, and the resulting figures are shown in (**Fig. S1**).

### Mapping bacterial species on the tree of life and detection of distance from LUCA

To illustrate evolutionary variations in ribosomal proteins on the tree of life, we used the tree of life from Ref. ^51^ as the scaffold that was uploaded to the interactive tree of life website (https://itol.embl.de/) where certain bacterial branches were highlighted to measure bacterial distances from the root of three of life or LUCA and then to create the illustration in (**Fig. 3**).

## Acknowledgements

We thank members of the Melnikov, Hirt, Schneider and Kapralov laboratories (Newcastle University, UK) for their feedback and help with preparing this manuscript. This work was supported by the NUORS award from Newcastle University (to K.H-B.), the BBSRC UK (4-year PhD studentship BB/T008695/1. to C.R.B.) and the Royal Society UK (RGS\R2\202003 to S.M.).

