## Supplementary Information for "Ribosomes as molecular thermometers: metal-binding sites in ribosomal proteins are robust indicators of bacterial adaptation to heat and cold"

### Content

#### **Supplementary Tables and Figures** (in this file)

1. **Supplementary Table S1 |**
   Databases used to collect values of optimal growth temperatures for bacterial species.

2. **Supplementary Table S2 |**
   The occurrence of metal-coordinating ribosomal proteins depends on the optimal growth temperature of bacterial species.

3. **Supplementary Figure S1 |**
   Comparison of consensus sequences for ribosomal proteins from heat-adapted species vs cold-adapted species.

#### **Supplementary Data** (attached as separate files)

1. **Supplementary Data 1 |**
   The list of bacteria and their optimal growth temperatures that were analyzed in this study.

2. **Supplementary Data 2 |**
   Sequences of ribosomal proteins analyzed in this study.

3. **Supplementary Data 3 |**
   Aligned sequences of ribosomal proteins analyzed in this study.

**Table S1 ǀ** Websites used for web scraping to collect information about the optimal growth temperatures of microbial organisms.

| Database | Database Full Name | URL |
|---|---|---|
| **ARS (NRRL)** | Agricultural Research Service Culture Collection, National Center for Agricultural Utilization Research, USA | https://nrrl.ncaur.usda.gov/ |
| **ATCC** | American Type Culture Collection, USA | https://www.atcc.org/ |
| **BCCM** | Belgian Coordinated Collections of Microorganisms, Belgium | https://bccm.belspo.be/ |
| **BCRC** | Bioresources Collection and Research Center, Taiwan | https://catalog.bcrc.firdi.org.tw |
| **CABRI** | Common Access to Biological Resources and Information | http://www.cabri.org/ |
| **CCARM** | Culture Collection of Antibiotics Resistant Microbes, UK | https://www.phe-culturecollections.org.uk |
| **CCM** | Czech Collection of Microorganisms, the Czech Republic | https://www.sci.muni.cz/ccm/ |
| **CCUG** | Culture Collection, University of Goteborg, Sweden | https://www.ccug.se/ |
| **CECT** | Spanish Type Culture Collection, Spain | https://www.uv.es/cect/ |
| **CICC** | China Center of Industrial Culture Collection | http://english.china-cicc.org |
| **CIP** | Center for Biological Resources of the Institute Pasteur, France | https://catalogue-crbip.pasteur.fr |
| **DSMZ** | German Collection of Microorganisms and Cell Cultures GmbH, Germany | https://www.dsmz.de |
| **GRIN** | Agricultural Genetic Resources Information Center, USA | https://www.ars-grin.gov |
| **HAMBI** | Culture Collection of Department of Applied Chemistry and Microbiology, University of Helsinki, Finland | https://kotka.luomus.fi/culture/bac |
| **JCM** | Japan Collection of Microorganisms (RIKEN Bioresource Center), Japan | https://jcm.brc.riken.jp/en/ordering_e |
| **KEMB** | Korea Environmental Microorganisms Bank, South Korea | https://kemb.or.kr |
| **NBIMCC** | National Bank for Industrial Microorganisms and Cell Cultures, Bulgaria | https://www.nbimcc.org/en/about.htm |
| **NBRC** | NITE Biological Resource Center, Japan | https://www.nite.go.jp |
| **NCIMB** | National Collections of Industrial, Food and Marine Bacteria, UK | https://www.ncimb.com/ |
| **NCMA** | National Center for Marine Algae and Microbiota (NCMA), USA | https://ncma.bigelow.org/ |
| **NCTC** | National Collection of Type Cultures, UK | https://www.phe-culturecollections.org.uk/ |
| **NIES** | National Institute for Environmental Studies, Japan | https://mcc.nies.go.jp/ |
| **VKM** | All-Russian Collection of Microorganisms, Russia | http://www.vkm.ru |

**Table S2 |** Websites used for web scraping to collect information about the optimal growth temperatures of microbial organisms.

| **Bacteria lineage** | **Average optimal growth temperature** | **Average number of metal-coordinating r-protein** |
|---|---|---|
| Acidobacteria | 23.3 | 7.3 |
| PVC group;Chlamydiae | 25 | 0.5 |
| Terrabacteria group;Cyanobacteria/Melainabacteria group | 26 | 2 |
| PVC group;Planctomycetes | 27.4 | 4.4 |
| Proteobacteria;Alphaproteobacteria | 27.7 | 0.4 |
| Chrysiogenetes;Chrysiogenetes | 28 | 7 |
| PVC group;Kiritimatiellaeota | 28 | 6 |
| Proteobacteria;Gammaproteobacteria | 28.6 | 1.5 |
| Proteobacteria;Betaproteobacteria | 28.9 | 1.1 |
| FCB group;Bacteroidetes/Chlorobi group | 29.4 | 0.5 |
| PVC group;Verrucomicrobia | 30 | 0.6 |
| Elusimicrobia;Endomicrobia | 30 | 6 |
| FCB group;Gemmatimonadetes | 30 | 5 |
| Proteobacteria;Oligoflexia | 30 | 2 |
| Terrabacteria group;Actinobacteria | 31.7 | 4.9 |
| Proteobacteria;delta/epsilon subdivisions | 31.9 | 6 |
| Terrabacteria group;Tenericutes | 33.5 | 4.2 |
| Fusobacteriia | 35.2 | 2.5 |
| Spirochaetia | 36.4 | 4.8 |
| Terrabacteria group;Firmicutes | 40.7 | 5.6 |
| Synergistia | 47.8 | 7.4 |
| Deferribacteres | 48 | 8 |
| Nitrospira | 48.5 | 5.5 |
| Terrabacteria group;Chloroflexi | 51.5 | 5.8 |
| Calditrichaeota;Calditrichae | 60 | 6 |
| Terrabacteria group;Coprothermobacterota | 60 | 6 |
| Terrabacteria group;Deinococcus-Thermus | 60.7 | 5.2 |
| Thermotogae | 64.6 | 7.5 |
| Caldisericia | 65 | 7 |
| Thermodesulfobacteria | 71.6 | 7.6 |
| Aquificae | 72.2 | 6.5 |
| Dictyoglomia | 75 | 8 |

**Figure S1 |** This figure compares two consensus sequences of each metal-coordinating ribosomal protein, with one of these consensus sequences corresponding to cold-adapted bacteria (on the left) and another one corresponding to heat-adapted bacteria (on the right).

**uL24**

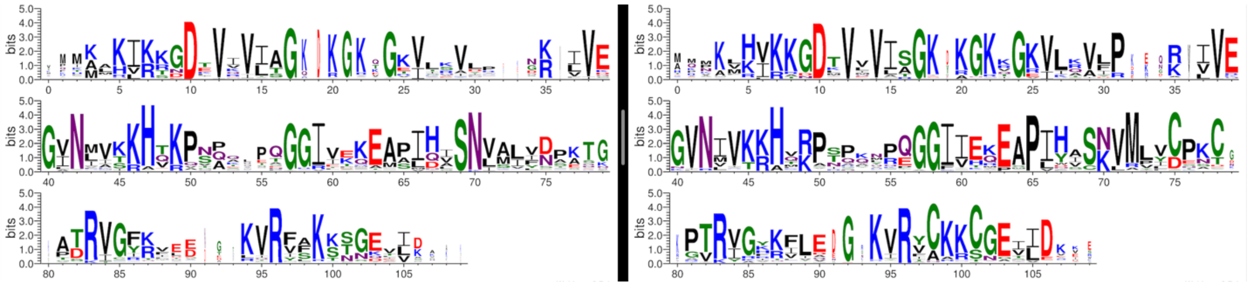

**bL28**

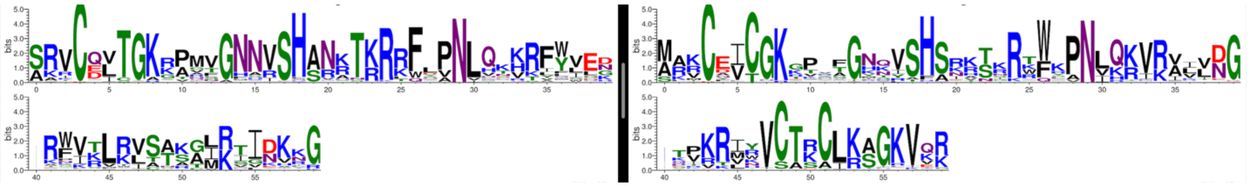

**bL31**

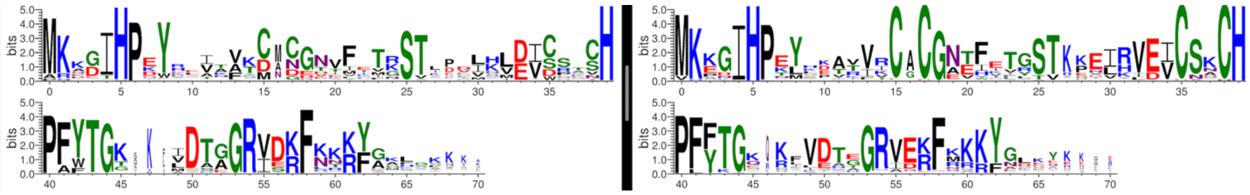

**bL32**

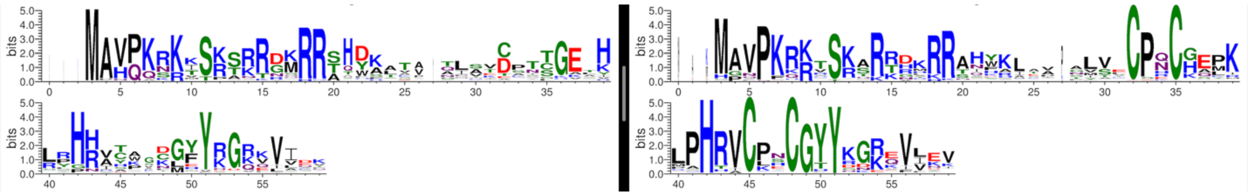

**bL33**

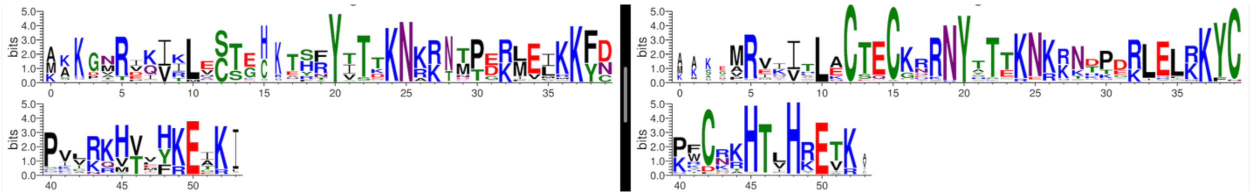

## bL36

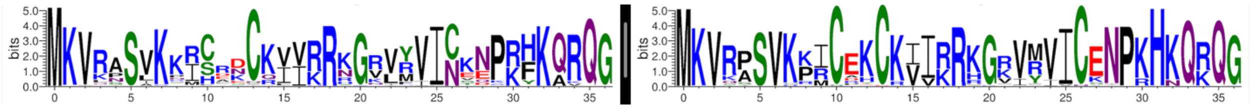

## uS4

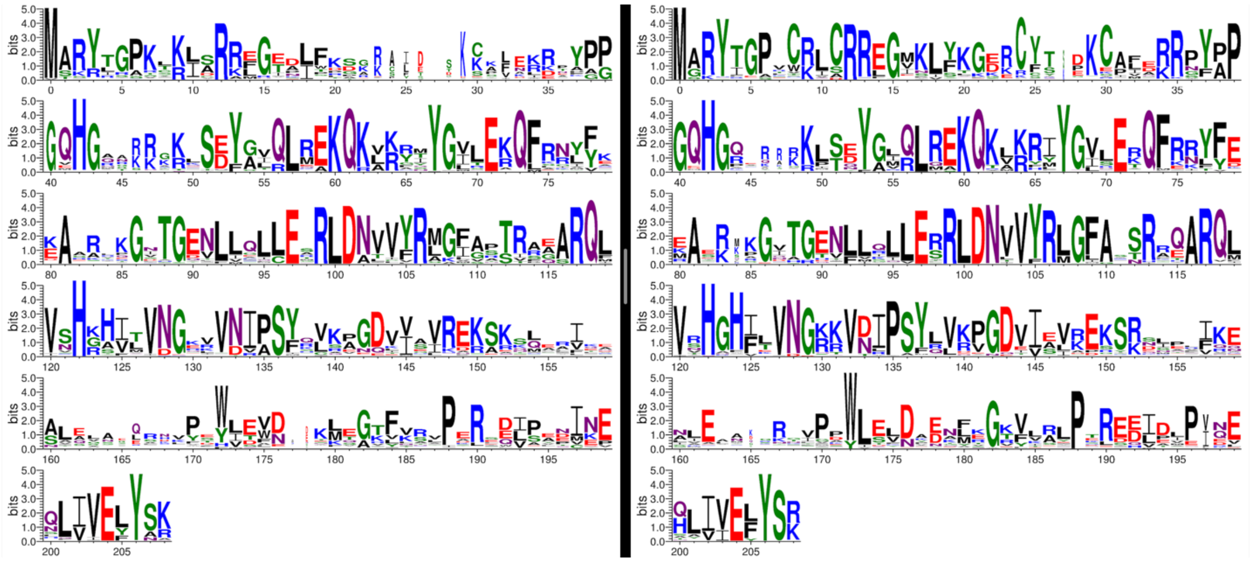

## bS18

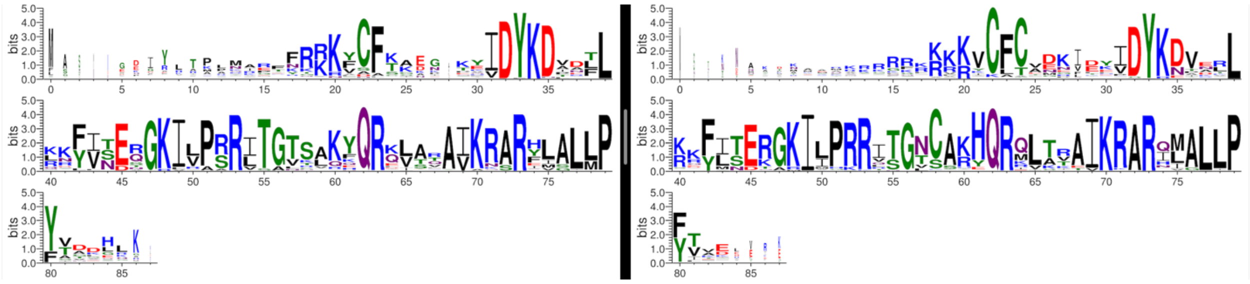
